# MobiSeq: De Novo SNP discovery in model and non-model species through sequencing the flanking region of transposable elements

**DOI:** 10.1101/349290

**Authors:** Alba Rey-Iglesia, Shyam Gopalakrishan, Christian Carøe, David E. Alquezar-Planas, Anne Ahlmann Nielsen, Timo Röder, Lene Bruhn Pedersen, Christina Næsborg-Nielsen, Mikkel-Holger S. Sinding, Martin Fredensborf Rath, Zhipeng Li, Bent Petersen, M. Thomas P. Gilbert, Michael Bunce, Tobias Mourier, Anders Johannes Hansen

## Abstract

In recent years, the availability of reduced representation library (RRL) methods has catalysed an expansion of genome-scale studies to characterize both model and non-model organisms. Most of these methods rely on the use of restriction enzymes to obtain DNA sequences at a genome-wide level. These approaches have been widely used to sequence thousands of markers across individuals for many organisms at a reasonable cost, revolutionizing the field of population genomics. However, there are still some limitations associated with these methods, in particular, the high molecular weight DNA required as starting material, the reduced number of common loci among investigated samples, and the short length of the sequenced site-associated DNA. Here, we present MobiSeq, a RRL protocol exploiting simple laboratory techniques, that generates genomic data based on PCR targeted-enrichment of transposable elements and the sequencing of the associated flanking region. We validate its performance across 103 DNA extracts derived from three mammalian species: grey wolf (*Canis lupus*), red deer complex (*Cervus sp*.), and brown rat (*Rattus norvegicus*). MobiSeq enables the sequencing of hundreds of thousands loci across the genome, and performs SNP discovery with relatively low rates of clonality. Given the ease and flexibility of MobiSeq protocol, the method has the potential to be implemented for marker discovery and population genomics across a wide range of organisms – enabling the exploration of diverse evolutionary and conservation questions.

## Introduction

Next generation sequencing (NGS) has revolutionized the world of genomics, allowing genome scale studies in model and non-model organisms (e.g. Davey et al., 2011; Ellegren, 2014; Goodwin, McPherson, & McCombie, 2016). Despite ongoing cost reductions in both the sequencing of reference genomes, and the resequencing of genomes from individuals and populations (Fuentes-Pardo & Ruzzante, 2017; Goodwin et al., 2016), it still remains cost-prohibitive for most research groups to generate whole genomes. Also, many research questions can be answered using a smaller set of SNPs that are measured in a subset of genomic regions (Davey et al., 2011). Thus, reduced representation library (RRL) methods have become a popular alternative for SNP discovery and genotyping (Davey & Blaxter, 2011), in particular for non-model organisms. Several RRL strategies have been developed in the last years, including restriction site-associated DNA sequencing (RADseq) (Baird et al., 2008; Davey & Blaxter, 2011), double digest RADseq (ddRADseq) (Peterson, Weber, Kay, Fisher, & Hoekstram, 2012) or genotyping-by-sequencing (GBS) (Elshire et al., 2011), as well as the combination of RRLs with hybridization by capture for genotyping museum and ancient specimens (Sánchez Barreiro et al., 2016; Schmid et al., 2017).

The core feature of traditional RRL techniques is the use of restriction enzymes to obtain DNA sequence at a genome-wide set of loci (Andrews, Good, Miller, Luikart, & Hohenlohe, 2016; Davey & Blaxter, 2011). The sequencing of a subset of the genome by RRLs provides a high depth of coverage per locus at a reduced cost (Andrews et al., 2016), and to date RRLs have been successfully used in population genomic (e.g. Hohenlohe et al., 2010), phylogeographic (e.g. Emerson et al., 2010; Gaither et al., 2015), and phylogenomic studies (e.g. Wagner et al., 2013). Despite all the advantages that traditional RRL methodologies present, some important challenges remain: (1) high molecular weight DNA is required for enzymatic digestion, (2) allele dropout leads to high proportion of missing data, (3) high percentage of clonal reads, and (4) relatively complex laboratory workflows. Hybridization capture of RRL loci circumvents some of these challenges, allowing the sequencing of RRL loci in degraded samples and reducing allele dropout. However, bait design can be complex and costly, and typically requires commercially synthesized oligonucleotide probes (Faircloth et al., 2012; Sánchez Barreiro et al., 2016; Schmid et al., 2017). Thus, there is a niche for methods that allow some flexibility in the initial DNA quality requirements, while simplifying laboratory workflows, and reducing costs. In this study we present MobiSeq, a novel NGS genotyping method that takes advantage of the conserved sequences of transposable elements (TE) as anchoring points in order to generate sequence data containing the TE sequence, as well as the genomic regions flanking the element, which can be used for genotyping and SNP discovery. The method has been developed with the focus on samples stored or preserved in sub-optimal conditions that yield average DNA fragments between 350 – 800 bp.

TEs are self-replicating mobile elements that insert themselves in new places of the genome, either through a cut-and-paste or a copy-and-paste mechanism (Kazazian, 2004). The latter, also referred to as retrotransposable elements or type II elements, are predominant in mammalian genomes (with bats as a notable exception (Ray et al., 2008)). LINE elements are long retrotransposable elements encoding the enzymatic machinery required for their own movement. In contrast, SINEs are short transcribed sequences – often derived from small RNA genes – that do not encode any proteins, instead relying on proteins encoded by LINE elements (Dewannieux, Esnault, & Heidmann, 2003). Insertions of LINEs and SINEs take place through the so-called target-primed reverse transcription, ensuring that the 3’ end of the elements are always present whereas the 5’ may be truncated (Luan, Korman, Jakubczak, & Eickbush, 1993). Most TEs display little insertion preference and can be scattered throughout the genome, although they are negatively selected in exonic regions (Sela, Mersch, Hotz-Wagenblatt, & Ast, 2010). They are found in almost all investigated eukaryotic genomes (Chénais, Caruso, Hiard, & Casse, 2012) and typically constitute more than 50% of the genome in mammals (e.g. International Human Genome Sequencing Consortium, 2001; Sotero-Caio, Platt, Suh, & Ray, 2016). The human and mouse genomes are by far the most well-studied animal genomes with regards to TE activity. Studies on these species have shown that the ongoing TE activity has resulted in genomic differences between closely related species (Mills et al., 2006; Yohn et al., 2005), populations (Akagi, Li, Stephens, Volfovsky, & Simer, 2008; Cordaux, Srikanta, Lee, Stoneking, & Batzer, 2007), and individuals (Konkel, Wang, Liang, & Batzer, 2007; Levy et al., 2007).

Several studies have used NGS to investigate TEs, especially those associated with humans and diseases (e.g. Ewing & Kazazian, 2010; Ewing & Kazazian, 2011, Tang et al., 2016; Tubio et al., 2014,). The MobiSeq method presented here can generate many thousands of unique sequences per sample, each containing the TE sequence and the flanking genomic regions, which are subsequently used for SNP discovery and genotyping. MobiSeq relies on a modified version of the blunt-end single-tube double-stranded DNA library construction protocol described in Carøe et al. (2017), coupled with a TE-target enrichment PCR step prior to sequencing. TE-target primers can be designed to enrich for any TE element present in the species of interest, making it a very flexible protocol for use on eukaryotic genomic DNA (Figure 1). Furthermore, several TEs can be combined, in order to increase the number of sequenced markers, thus increase the proportion of genome coverage and analytical resolution.

**Figure 1.**
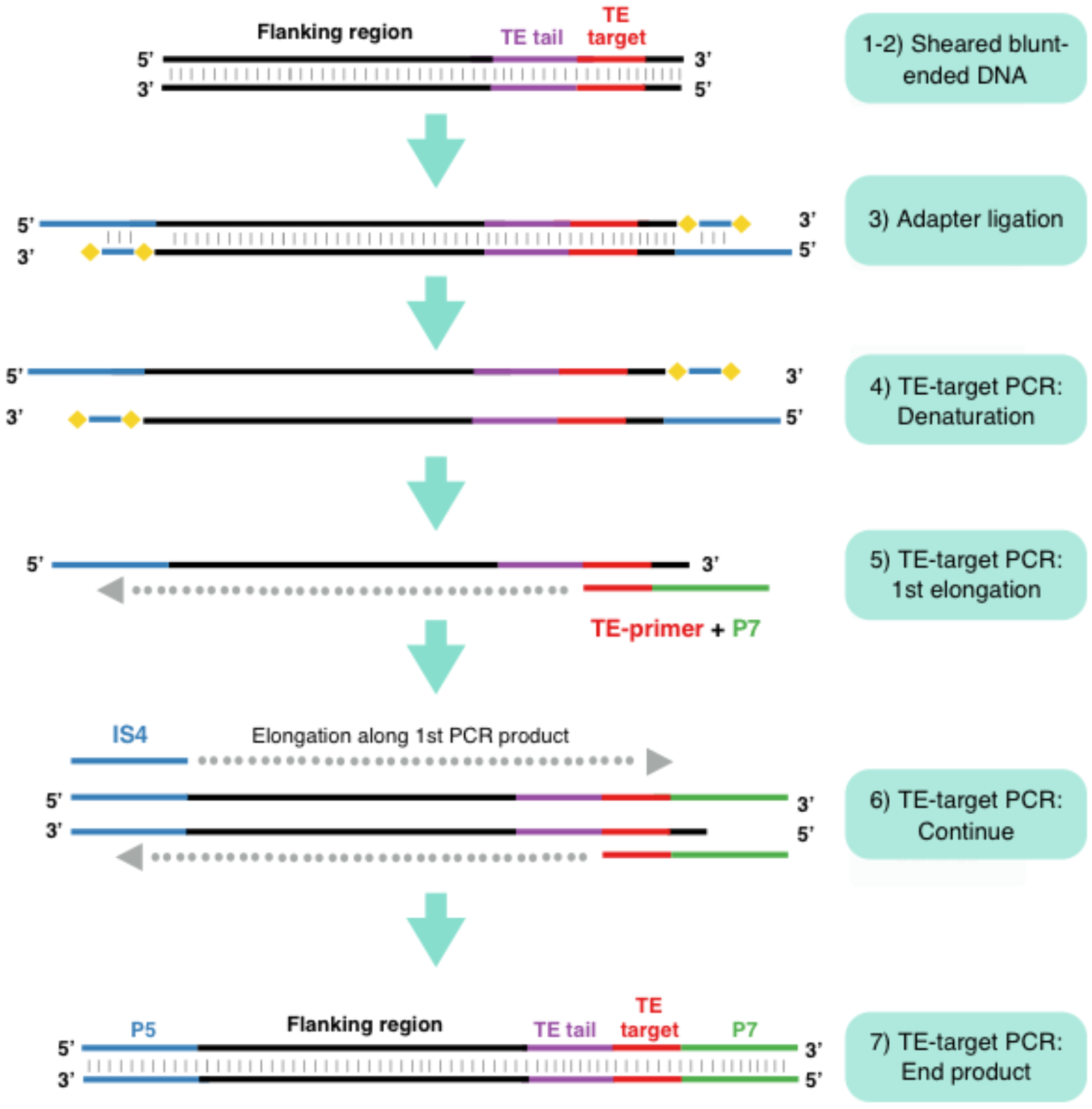
Schematic overview of the protocol. 1) Sample DNA is sheared (if required) into 600 – 800 bp average size fragments. 2) During blunt-end repair, overhanging 5’ and 3’ ends are filled in or removed by T4 DNA polymerase. 5’ phosphates are attached using T4 polynucleotide kinase. 3) Double-stranded mP5 adapters are ligated to the fragment by T4 DNA Ligase. Adapters do not carry 5’ phosphates and therefore only one strand is ligated to the fragments. 4) TE-target PCR is performed using forward IS4 primer and the TE-fusion primer. TE-target PCR will also result in the incorporation of P7 sequencing adapter. 5) PCR elongation will only occur upstream of the 3’ end of the TE-target primer. 6) The end product of the TE-target PCR will be the TE-target sequence, a TE tail and the flanking genomic region. It will also contain Illumina sequencing adapters (P5 and P7). This PCR product can be indexed using single or dual indexing primers, then pooled for sequencing with other samples. Yellow diamonds represent mP5 adapter modifications.

To demonstrate efficacy, we report MobiSeq data on three mammal species: brown rat (*Rattus norvegicus*), grey wolf (*Canis lupus lupus*), and red deer complex species (*Cervus sp*). Two fallow deer samples (*Dama dama*) were also included to test the usefulness of the method when working with distantly related species (e.g. red deer and fallow deer). Thus, we demonstrate its applicability and flexibility for genotyping model and non-model organisms, and discuss both possible future applications in the field of population genomics, and potential limitations.

## Materials and methods

### Sample information

DNA was extracted from 103 brown rat, grey wolf, and deer samples, see Table S1 (Supporting Information) for a detailed description of the specimens included in the study. The deer samples used in the study were a mix of modern (1996 – 2015) and historic material (collected between 1909 – 1960). The rat sample set consisted of four male Sprague-Dawley rat individuals (Charles River Laboratories, Germany), each sampled from 13 different tissue types. Species selection was based on three criteria: 1) a model species with a well-characterized genome and TE variation in the genome (brown rat); 2) a species with good quality genome assembly available and studied extensively in population genetic studies (grey wolf) (e.g. Kardos et al., 2018; Pilot et al., 2014; Rutledge, Devillard, Boone, Hohenloge, & White, 2015); and 3) a less well-studied complex of species (i.e. the red deer complex). DNA extractions were performed using DNeasy Blood and Tissue (Qiagen) following manufacturer’s instructions. DNA elution was performed twice in 50 μl AE buffer and with 10 minutes of incubation time at 37 °C prior to elution, in order to increase DNA yield. Extractions were quantified using a Qubit dsDNA High Sensitivity (HS) assay (Life technologies). Samples with higher DNA concentration than HS assay range were quantified again with a Qubit Broad Range assay (Life technologies).

### TE-target primer design

In order to enrich our libraries for specific TE-target elements, we designed four TE primers for repeat families that showed hallmarks of recent activity. From the RepeatMasker (Smit, Hubley, & Green, 2013) annotations of the rat and the dog genome (Speir et al., 2015; Rat Genome Sequencing Consortium, 2014; Lindblad-Toh et al., 2005), we extracted 3’ tails from TE loci that were nearly full-length and showed low levels of divergence from the consensus sequence. Sequences were extracted using samtools (Li et al., 2009) and aligned with muscle (Edgar, 2004). Highly conserved regions were determined from the alignments, and selected as potential primer sites. This resulted in one primer targeting a L1 TE (a superfamily of LINEs) in brown rats (L1RnT1: 5’ CCGGAAACCGGGAAAGGGAATAACAC 3’); and two primers targeting two different TE in grey wolves (SINE: 5’GAGACCCGGGATCGAATCCC 3’; LINE: 5’ GATAGCCAAACTGTGGAAGG 3’). Due to the lack of a well-annotated genome for the red deer complex, a SINE primer was selected from the study Nilsson, Klassert, Bertelsen, Hallström and Janke (2012); (BOV2A: 5’ GGGACGGGGGAGCCTGGTGGGCTG 3’). The TE-target oligonucleotides were then combined with the sequence of the P7 adapter (Meyer & Kircher, 2010), in order to create a fusion primer (TE+P7) that would enrich for TE sites, at the same time as adding the P7 sequencing adapter adapter compatible with binding to Illumina flow cells (see Table S2 for an overview of the oligonucleotides).

### Modified P5 adapter

A modified version of the P5 adapter from Meyer and Kircher (2010) was designed for this protocol. In this modified P5 (mp5), the IS1 oligo is kept as in Meyer and Kircher (2010). However, IS3 oligonucleotide presents a modified sequence by adding a C3 spacer at the 3’ end blocking polymerase extension. This, together with the conventional lack of a 5’-phosphate, allows us to run a PCR reaction using a universal primer for the adapter sequence (IS4 or IS7, see Table S2, Supporting Information) and a TE-target primer enriching for a specific subset of TEs (Figure 1). Hybridization of IS1 and the modified IS3 to generate mp5 was performed as in Meyer and Kircher (2010).

### Methods for preparation of sequencing libraries

An overview of the method is represented in Figure 1 (full protocol available as Appendix S1, Supporting information). Prior to library build, DNA extracts were fragmented using Bioruptor NGS (Diagenode) to an average length of 600 – 700 bp. Four of the samples were not fragmented prior to library build, as their average DNA fragment size was ca. 350 – 400 bp (Table S1, Supporting information). Starting material for Bioruptor varied across samples (between 200 – 2000 ng), depending on the DNA extract concentration (Table S1, Supporting information). Biorupted DNA was size selected using magnetic beads purification, either Agencourt AMPure XP (Beckman Coulter) or Sera-Mag Speedbeads (ThermoScientific) at 0.7x to remove DNA fragments shorter than ca. 200 bp. Size selected DNA was then quantified using Qubit HS reagents, as previously described, and loaded in a TapeStation High Sensitivity (TS-HS) 2200 (Agilent) to obtain a length profile of the fragmented material. Illumina library building was based on a recently developed blunt-end single-tube protocol (Carøe et al., 2017) with some modifications as in Mak et al. (2017) (full protocol available as Appendix S1, Supporting information). In particular, the ligated adapters differed from the ones in the original protocol by excluding the use of a P7 adapter (only using the mp5 adapter) and excluding the adapter fill-in reaction. Following library preparation, the reactions were purified using a magnetic beads purification protocol, Agencourt AMPure XP (Beckman Coulter) or Sera-Mag Speedbeads (ThermoScientific), at 0.70x. Purified libraries were eluted in 30 μl of EBT.

### TE-enrichment PCR

Libraries were enriched for fragments containing the TE of interest by using a TE-enrichment PCR. Primers for this PCR were forward primer IS4 or IS7 (Meyer & Kircher, 2010) and the fusion reverse primer described in prior sections (Table S2 for oligonucleotide sequences, Supporting information), see Figure 1 (TE-enrichment PCR). PCR reactions were performed in 25 μl containing: 1x Accuprime Pfx Mix (ThermoScientific), 0.4 μM of each primer, 2% DMSO (ThermoScientific), 0.02 U/μl Accuprime Pfx polymerase, and 5 μl of the purified libraries. Cycling parameters were denaturation at 95 °C for 30 seconds, followed by 10-15 cycles of denaturation at 95 °C for 30 seconds, annealing between 60-67 °C (depending on the TE-target primer) 30 seconds, and extension at 68 °C for 60 seconds). TE-enriched libraries were purified using magnetic beads at 1x. Purified DNA was eluted in 30 μl of EBT. Concentration was measured using a Qubit HS assay and loaded in a TS-HS 2200 (Agilent).

### Index PCR and sequencing

TE-enriched libraries were indexed and amplified for sequencing as described in Meyer and Kircher (2010). PCR reactions were performed in 25 μl containing: 1x Accuprime Pfx Mix (Thermo Fisher), 0.4 μM of each primer, 2% DMSO, 0.02 U/μl of the enzyme, 5 μl of the purified libraries. Cycling parameters were denaturation at 95 °C for 30 seconds, followed by 5-10 cycles of (denaturation at 95 °C for 30 seconds, annealing at 60 °C for 30 seconds, and extension at 68 °C for 60 seconds). Indexed libraries were purified using magnetic beads at 1x. Purified DNA was eluted in 30 μl of EBT. Concentration was measured using a Qubit HS assay and loaded in a TS-HS 2200 (Agilent). Indexed libraries were pooled and sequenced at the Danish National High-throughput Sequencing Centre, Copenhagen, Denmark, on an Illumina MiSeq Instrument for 250 cycles in paired-end read mode. The sequencing architecture is illustrated in Figure 2.

**Figure 2.**
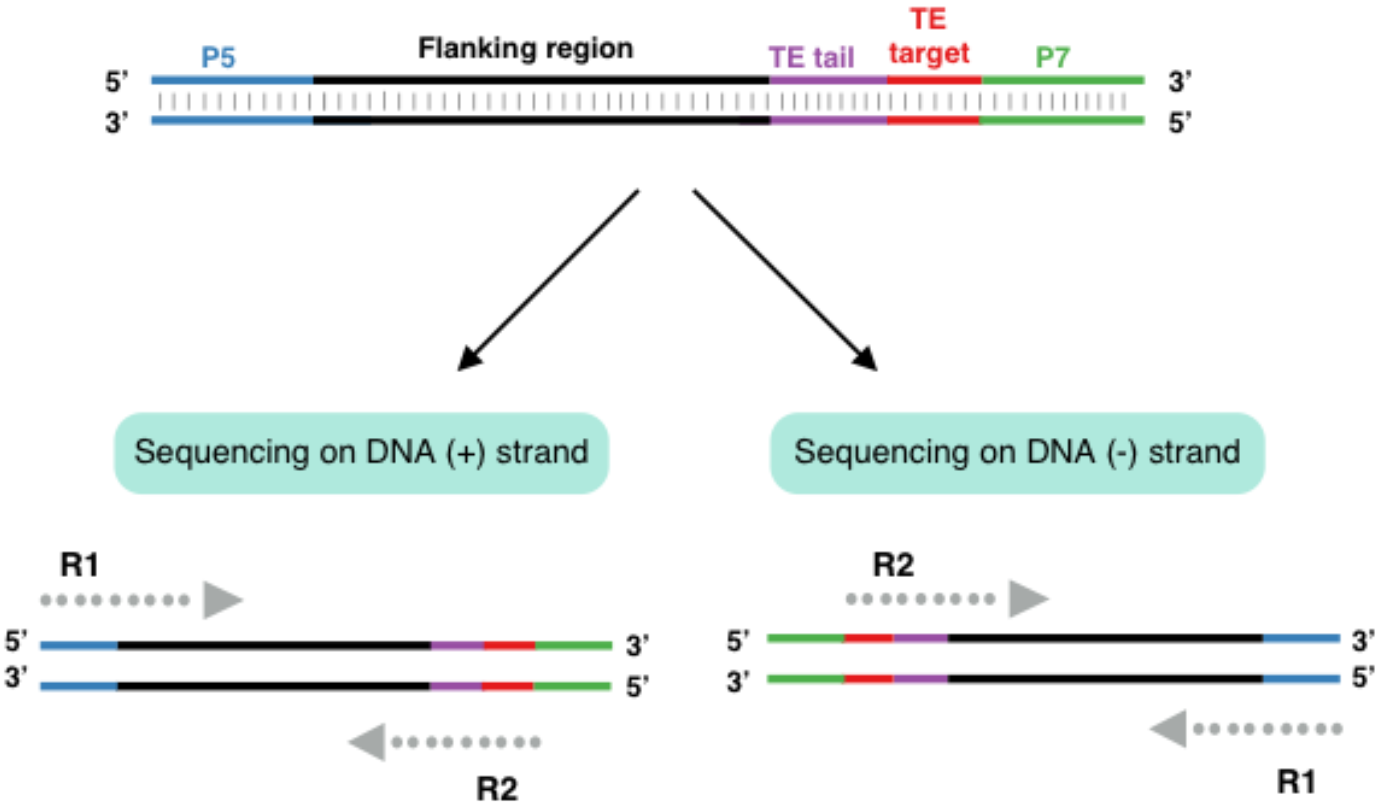
Sequencing architecture of Transposable Element (TE) flanking region in the (+) and (−) strand. The TE-target primer will always be at the 5’end of Read 2 (R2). Sequencing into the flanking will only happen from the 3’ end into the flanking region. The start of Read 1 (R1) will be variable, as DNA was random sheared, which allows the identification of PCR duplicates.

### Data processing

Raw reads were filtered based on the presence of the TE-target primer sequence in the 5’end of the reverse R2 (Figure 2, Supplementary Figure 1). Reads without the TE-target primer sequence were discarded, as they represent off-target amplifications or PCR artefacts. Reads derived from the rat samples were merged by individual (see Table S1, Supporting information) and treated as such in all downstream analyses. Paired-end filtered reads were trimmed of adapter sequences, reads shorter than 25 bp were discarded and quality filtered using PALEOMIX (Schubert et al., 2014). Retained paired-end reads were mapped against a reference genome of the different species included in the study: brown rat genome (GCA_000001895.4; Rat Genome Sequencing Consortium, 2014); wolf genome (Gopalakrishnan et al., 2017); and red deer genome (Zhipeng Li et al., unpublished). Mapping was performed with BWA (Li & Durbin, 2010) using the mem algorithm and soft-clipping. PCR duplicates and reads mapping to multiple genomic locations were marked using Mark Duplicates (https://broadinstitute.github.io/picard/). GATK (DePristo et al., 2011) was used to perform indel realignment. Basic sequencing statistics, such as read numbers and clonality were quantified within the PALEOMIX pipeline (Schubert et al., 2014). For all downstream analysis, only reads mapping to loci (TE sites) that occur in more than 90% of the samples were retained using the bedtools software suite (Quinlan & Hall, 2010). The pairwise coverage comparisons for the retained set of loci were estimated for each primer pair using samtools bedcov (Li et al., 2009).

Aggregate coverage plots were generated for each sample and dataset using agPlus (Maehara & Ohkawa, 2015) from the primer site up to 1 kb of the flanking region. The total coverage was used to correct for sequencing depth differences between samples. Forward and reverse reads derived from (+) and (−) strand sequencing were merged into one plot, as well as analysed separately to detect strand-associated biases (Figure 2). Average GC content (%) per loci was estimated across datasets to explore the variation of coverage in relation to GC content across loci. PRESEQ (Daley & Smith, 2013) was used to infer the complexity of each library for all the sequenced loci.

ANGSD (Korneliussen, Albrechtsen, & Nielsen, 2014) was used to perform SNP calling, requiring a minimum mapping quality of 30, minimum base quality of 20, minimum depth of coverage of 3x per genotype, and coverage in at least 50% of individuals. NgsDist (Vieira, Lassalle, Korneliussen, & Fumagalli, 2015) was used to estimate the pairwise distances between the samples for each dataset using the SNP data generated with ANGSD. Trees were estimated from pairwise genetic distances with 100 bootstrap replicates using FastME (Lefort et al., 2015) and RaxML (Stamatakis, 2014, https://github.com/amkozlov/raxml-ng).

## Results

### Number of sequenced loci, coverage, depth and clonality

Our sequencing yielded a total of 10,366,644 raw reads. An average of 95% of the reads across datasets contained the TE-target primer in R2. Less than 1% of the reads were discarded after trimming adapters and quality filtering, and more than 99% of the remaining reads were mapped to each specific genome. Between 24% and 0.4% of the reads were discarded after duplicate removal. One of the deer specimens presented a higher number of PCR duplicates than average (see Discussion). See Table 1 and Table S3 (Supporting information) for more detailed sequencing and mapping statistics.

**Table 1.** Average sequencing and mapping statistics per TE-target primer pair. More detailed sequencing and mapping statistics per sample can be found in Table S3 (Supporting Information).

| Average | Total sequenced reads | Reads with valid primer | Reads after adapter removal | Reads mapping | Reads mapping % | Reads mapping after duplicate removal | Read mapping % after duplicate removal | Clonality % |
| --- | --- | --- | --- | --- | --- | --- | --- | --- |
| **Wolf_LINE** | 431,916 | 405,378 | 405,262 | 405,011 | 99.95 | 351,119 | 99.99 | 11.7 |
| **Wolf_SINE** | 959,580 | 929,932 | 928,504 | 928,540 | 99.90 | 894,928 | 99.99 | 3.3 |
| **Deer_BOV2A** | 1,695,644 | 1,593,592 | 1,590,334 | 1,589,287 | 99.96 | 1,507,708 | 99.98 | 4.5 |
| **Rat_L1** | 7,279,504 | 6,941,756 | 6,941,686 | 6,930,739 | 99.84 | 6,840,096 | 99.99 | 1.3 |

Our sequencing strategy yielded a variable number of TE-enriched loci across genomes, depending on the TE-target primer used for the different species. Note that for number of loci estimations and downstream data analyses, BOV2A was divided in two datasets (1) BOV2A_all including all the cervid samples, and (2) BOV2A_CE including only *Cervus* sp. (Table S1, Supporting information). Table 2 summarizes all the TE-loci identified, as well as those found in at least 90% of the samples. In all subsequent downstream analyses, only the loci at the 90% cut-off were used.

**Table 2.** Total number of TE loci identified in our samples and number of loci identified in 90% of the samples within each dataset.

| TE-target | All loci | 90% cut-off loci |
| --- | --- | --- |
| **Wolf_LINE** | 33,761 | 9,472 |
| **Wolf_SINE** | 191,751 | 36,596 |
| **BOV2A_all** | 749,606 | 23,208 |
| **BOV2A_CE** | 653,088 | 24,134 |
| **Rat_L1** | 270,726 | 61,845 |

We performed pairwise comparisons of the coverage at primer sites across the 90% cut-off loci across the different datasets. Supplementary Figures 2 – 5 represent the 90% cutoff loci pairwise comparisons. Wolf_LINE and Wolf_SINE, and Rat_L1 showed high primer coverage correlation across samples, which suggests that sequencing across loci and samples was similar. BOV2A presented more variation in primer coverage, with a few samples showing very low pairwise correlations to the rest of the samples.

Aggregate coverage plots show that the highest coverage values are derived from the – 250 bp to 0 bp of the distance from the primer site (Figure 3). This region represents all the R2 reads carrying the TE-target primer sequence. The slight increase in coverage before 0 is derived from R1 reads that overlap with R2. This is correlated with DNA fragment length and the sequencing mode used in this study, MiSeq 250 PE (i.e. short DNA fragments will result in high amount of overlap between the two reads in the read pair). The shorter average size in the BOV2A material clearly reflects this (Figure 3). After position 0, that represents the primer site, there is a drop in coverage that will continue decreasing, as the sequencing moves into the flanking region (Figure 3). Depth of coverage was variable between datasets; in general, the highest coverage values were derived from the wolf LINE dataset. The recovered flanking region was variable between datasets and it is influenced by initial DNA fragment size. BOV2A aggregate plot serves as an example of this. Supplementary Figures 6 – 9 show aggregate coverage plots split by (+) and (−) strand.

**Figure 3.**
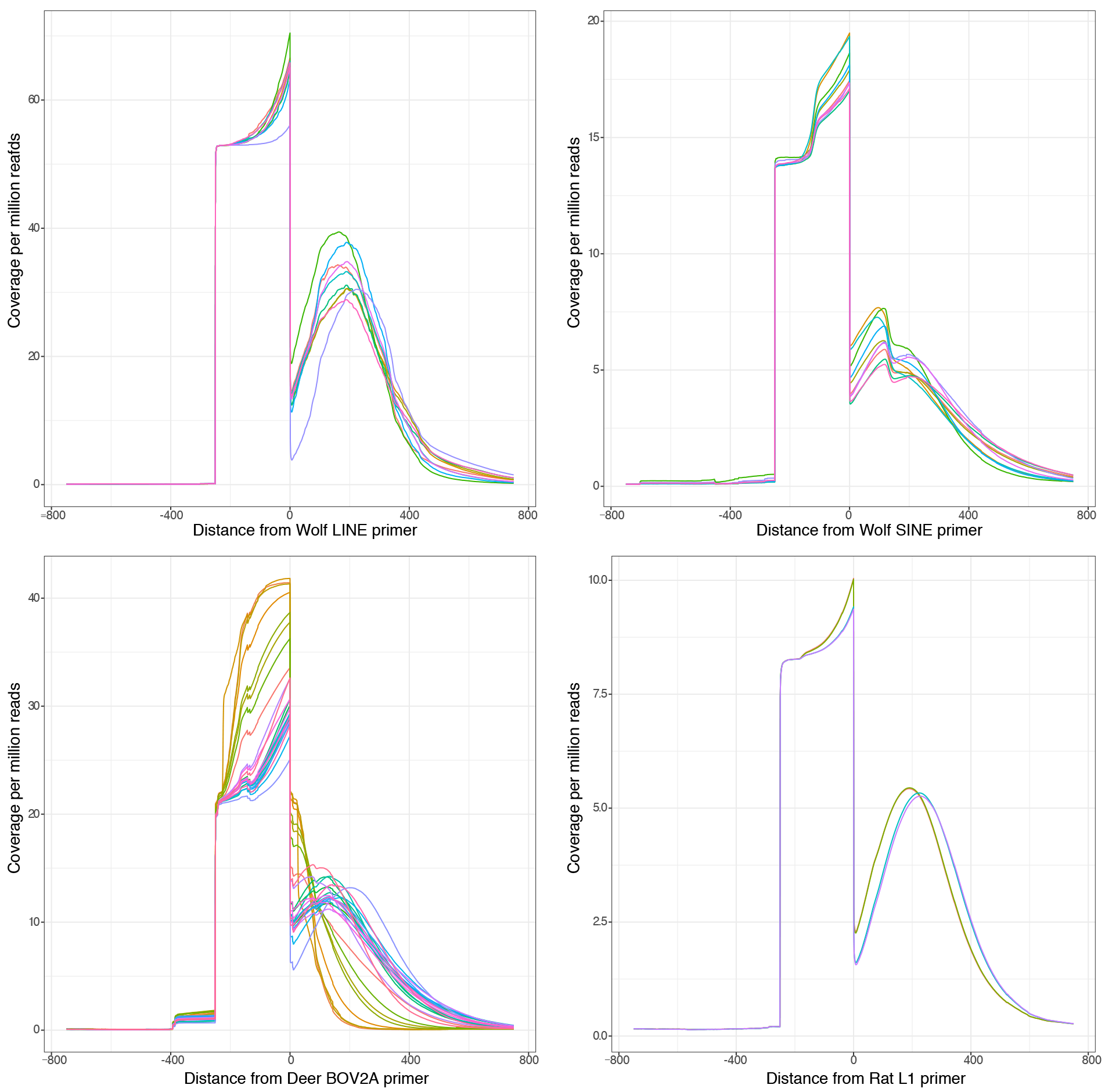
Aggregate coverage plots generated using agplus. Y-axis represents coverage per million reads; coverage was normalized to mitigate read number differences between samples. X-axis represents distance from TE-target primer. Zero represents the start of the TE-target primer. Each line represents a single individual.

### SNP discovery and genotyping

The sequenced TE-enriched loci yielded variable number of SNPs for the different TE-target primers. The total number of SNPs detected using ANGSD was: Wolf_LINE = 68,961; Wolf_SINE = 81,292; BOV2A_all = 210,949; BOV2A_CE = 114,973; and Rat_L1 = 64,482. We explored the variability in SNP numbers in relation to the minimum number of individuals sharing SNPs (Figure 4). The general trend is an increase in the number of shared SNPs as the number of individuals increases. However, we detected a drop in the number of shared SNPs for some of the primer pairs: Wolf_SINE (from 9 to 10 individuals), BOV2A_all (from 26 to 28 individuals), and BOV2A_CE (from 24 to 26 individuals). This drop seems to be associated with samples that performed poorly in the TE-enrichment PCR. We also investigated minor allele frequencies in the wolf and deer datasets (the rat dataset was excluded from these analyses because of the low number of samples) (Figure 4). The dataset BOV2A_CE presented a larger number of rare alleles than the other datasets.

**Figure 4.**
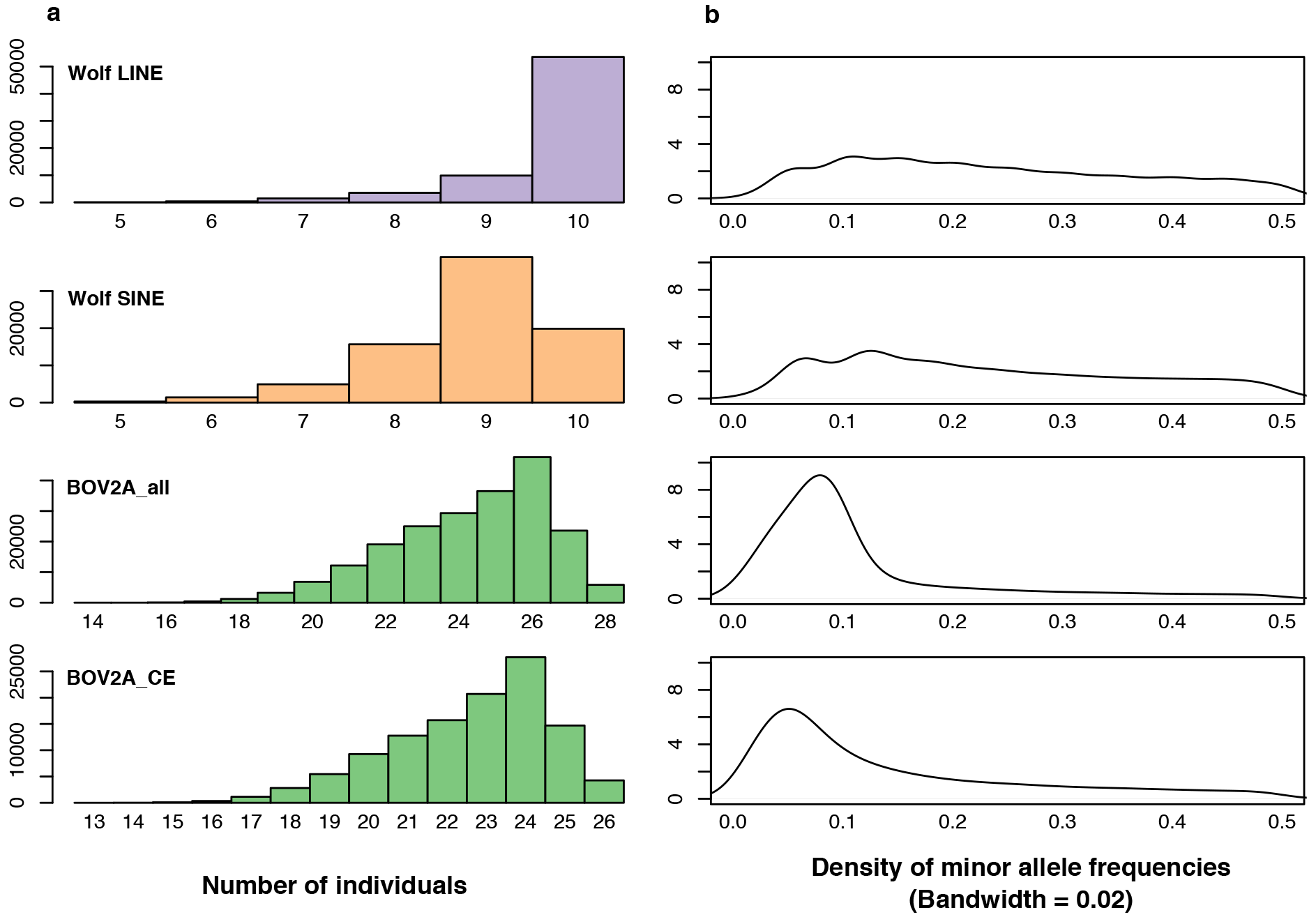
(**a**) Histogram of the number of SNPs (with coverage > 3) shared vs. the minimum number of individuals required in ANGSD. (**b**) Distribution of minor allele frequencies. The y-axis represents the density of minor allele frequencies.

We investigated cumulative rate of loci and SNPs sequenced per Megabase across genome scaffolds for each dataset (Supplementary Figure 10). Our results indicate that MobiSeq TE-target PCR amplified loci across all the scaffolds in the genome assembly, thus allowing us to perform random genone-wide SNP discovery across datasets. Finally, we explored SNP distribution with respect to the distance from the primer site (Supplementary Figure 11). Our results show that as sequencing moves into the flanking region, the number of discovered SNPs decreases. We note that the pattern of the number of SNPs discovered is reflective of the pattern of coverage across the loci (Figure 3). As an example of the applicability of MobiSeq data to estimate evolutionary relationships, we generated NGS distance trees based on SNPs for the wolf and deer datasets (Supplementary Figures 12 – 14).

### GC content and coverage

We explored the relationship between depth of coverage and GC content of the sequenced loci (Figure 5). Average GC content (%) in the sequenced loci is similar to the average GC content in the respective mammalian genomes. Except for the BOV2A_all dataset, the highest coverage is usually derived from the regions with around 45% GC content and it drops for windows with GC content more than 50% – 55%.

**Figure 5.**
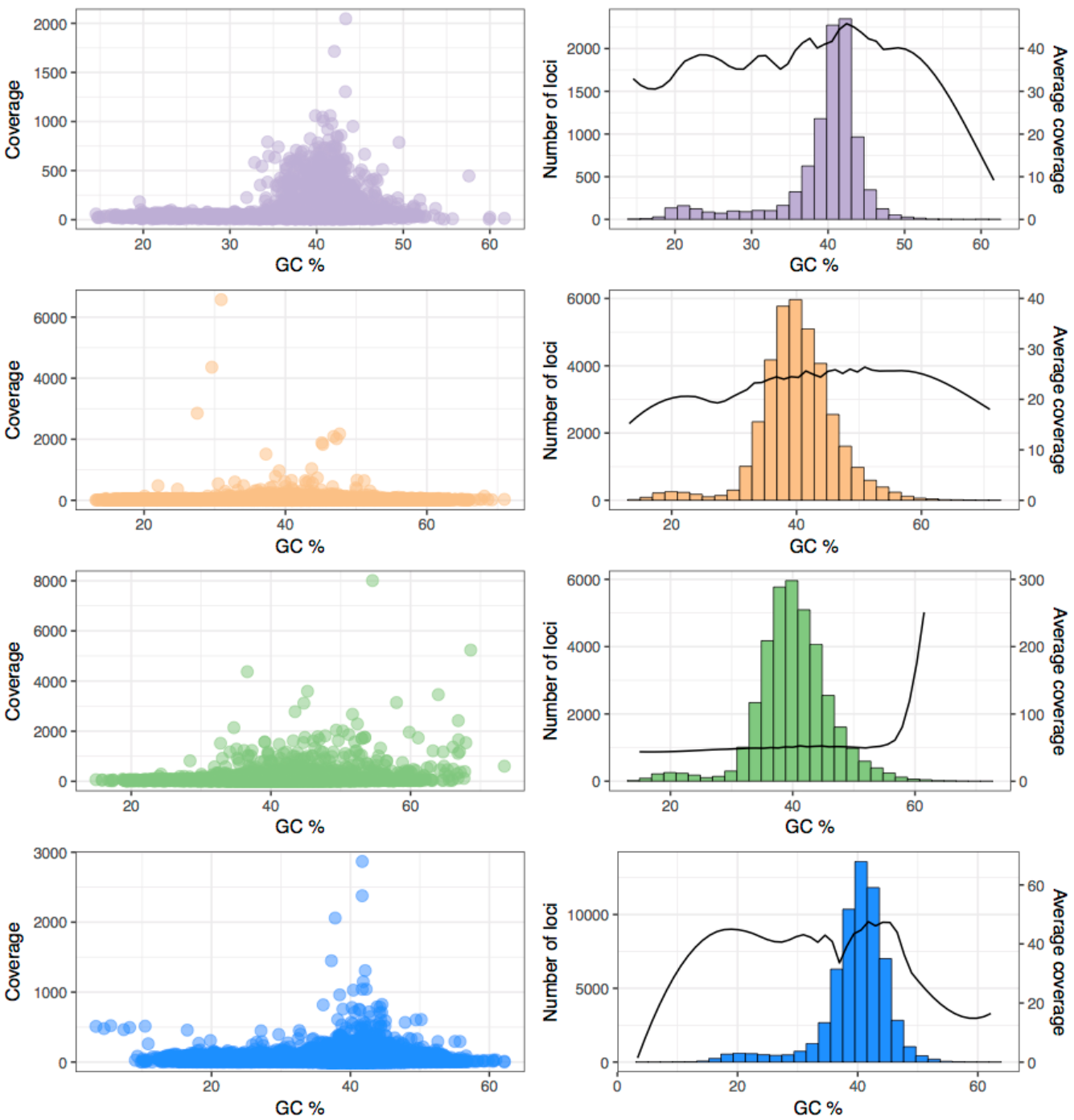
GC content (%) variation and coverage. Left panel represents average coverage expressed as average number of reads across loci versus GC (%). Right panel represents average GC content (%) in the different loci as a histogram, as well as the average coverage variation at those loci. Colours indicate the different datasets: lilac = Wolf_LINE; orange = Wolf_SINE; green = BOV2A_all; blue = Rat_L1.

### Library complexity

We used PRESEQ to estimate and predict the complexity of the MobiSeq libraries across all TE-target primers and samples (Figure 6). Our results indicate that MobiSeq libraries were sufficiently complex and that a sequencing plateau was not reached. Thus, further sequencing of these libraries would increase the number of detected loci and, as a result, the number of discovered SNPs that could be used to perform genomic inferences.

**Figure 6.**
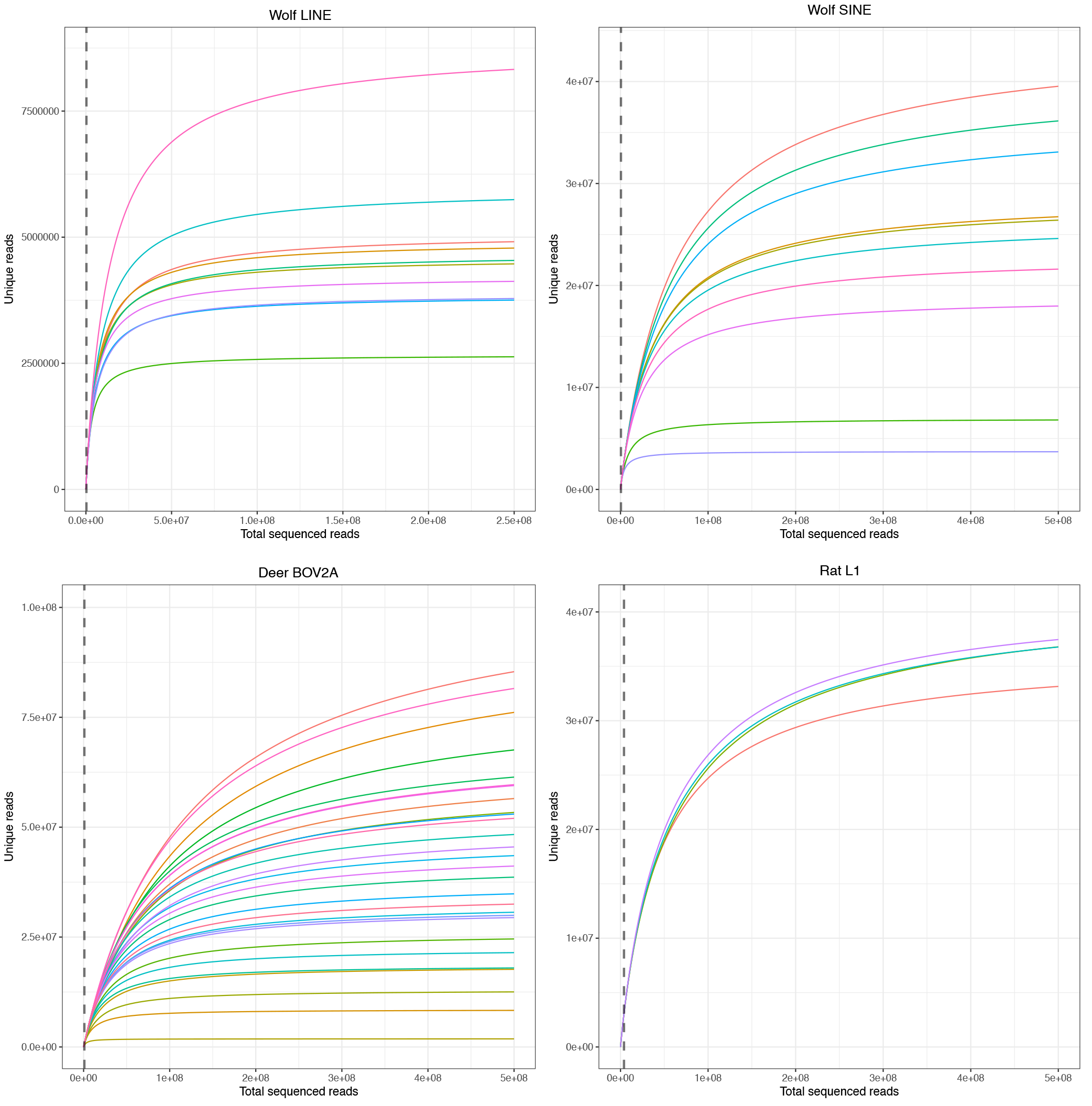
Plotted values generated by PRESEQ showing estimated number of distinct reads for each TE-target primer. Vertical dashed lines represent the average sequencing reads generated for each dataset.

## Discussion and conclusions

In this study, we have demonstrated the feasibility and adaptability of MobiSeq in three different taxa across a variety of tissues using both modern and degraded historical DNA.

### Amount of input DNA needed and Quality

Compared to other RRL protocols, the DNA quality and input requirements for MobiSeq offers a degree of flexibility. In general, traditional RRL methods require high molecular weight genomic DNA, which represents a limitation for poorly preserved samples (Andrews et al., 2016). In MobiSeq, highly fragmented material would result in fragmentation of the TE-target primer sites, which would reduce PCR success and affect the number of targeted loci. Also, the DNA quality of the starting material will have an influence on the length of the flanking region (i.e. highly fragmented DNA will reduce the sequenced flanking region). The historic deer specimens included in this study yielded an average DNA fragment size around 350 bp in the starting material (Table S1, Supporting information). Despite the underperformance of these samples in terms of clonality, coverage, and length of the flanking region, library complexity, and amount of missing data, we were still able to recover sufficient informative genomic data to include them in our NGS distance-based trees (Supplementary Figure 12). Nevertheless, whenever possible, we would suggest average DNA fragment sizes between 700 – 800 bp as, empirically, this is the material that has yielded the optimal results (i.e. number of sequenced SNPs and loci) in our analyses.

DNA input amount can be highly variable. We would suggest building the library on at least 100 ng of total input DNA. The current required DNA input in MobiSeq is similar to the amount used in RAD methods (Andrews et al., 2016). However, ongoing experiments indicate that starting material can be reduced down to 50 ng (Rey-Iglesia et al., unpublished).

### Duplicates

One of the limitations of RRL methodologies is the clonality generated due to PCR artefacts that would lead to increased artificial coverage and would affect SNP calling. For instance, several studies reported that PCR duplicates occur at high frequencies in RADseq data (e.g. Andrews & Luikart, 2014, Schweyen, Rozenberg, & Leese, 2014). PCR duplicates can be identified in RAD protocols that include a random shearing step and paired-end sequencing, like the original RADseq (Andrews et al., 2016; Davey et al., 2013). However, PCR duplicates cannot be identified in some of the other RAD strategies, because all fragments for a given locus will have identical start and stop positions (Andrews et al., 2016; Davey et al., 2011). Alternatives for controlling PCR duplicates are using PCR-free protocols, such as ezRAD that relies on Illumina PCR-free kits for library build. However, PCR-free methods are typically more expensive and require large amounts of starting material (Andrews et al., 2016).

In MobiSeq, the random shearing of the DNA prior to library build and the TE-target PCR set up generates fragments with different starting points in the 5’ start of the flanking region. This allows identification of putative PCR duplicates, based on the assumption that any read pair with identical starting position of the paired-end read are duplicates. However, in general, we obtained low values of clonality across datasets (Table 1, Table S3, Supporting information). Only one of the samples included in the study, a specimen from the BOV2A dataset, presented clonality values > 20%. The low clonality values obtained are associated with the high complexity of the libraries, due to the abundance of the targeted elements in the genome and the amplification success of our TE-target PCR approach.

### Depth of coverage

The highest depth of coverage was detected in the region closest to the primer site (ca. 250 bp representing the length of R2), coverage decreases as sequencing moves into the flanking region. Nevertheless, the average depth of coverage across the loci allowed for robust SNP calling using standard variant calling methods, as well as likelihood methods, such as those implemented in ANGSD.

Three principal factors that could be influencing the average depth of coverage are (i) high numbers of the targeted TEs, (ii) the sequencing strategy, and (iii) GC content of the sequenced loci. We sequenced our libraries using Illumina MiSeq 250 PE chemistry, obtaining an average number of reads per sample ranging from 431,916 in wolf_LINE to 1,695,644 in BOV2A. Higher numbers were recovered for rats (Table 1, Table S2, Supporting information), as each individual represents a pool of 13 libraries. A study on *Canis* admixture using RADSeq reported raw reads numbers between 8 – 37 million per sample (Rutledge et al., 2015). Other studies report average values of raw read numbers between (2 – 3 million reads) using RADSeq (e.g. Lah et al., 2015; Skovrind et al., 2016). Considering the total number of TE loci per dataset and the PRESEQ results, we would suggest using sequencing platforms with higher throughput (e.g. Illumina HiSeq 4000) to increase the amount of generated data, as well as depth of coverage. This would also allow us to obtain a more accurate comparison of MobiSeq with other RRL methodologies. The depth of coverage seems to also be influenced by the GC content of the sequenced loci. In general, our results show that regions containing around 45% GC content present the highest depth of coverage.

### Allele dropout

RRL datasets are likely to contain high proportions of missing data, mostly due to polymorphisms in the restriction enzyme recognition site (Gautier et al., 2013). In RRL methods, when a polymorphism occurs at a restriction enzyme recognition site, the enzyme will fail to cut the genomic DNA at that location, leading to allelic dropout (Andrews & Luikart, 2014; Andrews et al., 2016; Gautier et al., 2013;). Thus, even though high numbers of loci are sequenced per sample, the number of comparable sites can become highly reduced (Gautier et al., 2013). We believe that allelic dropout in MobiSeq is mostly dependent on DNA degradation, PCR biases, and distribution of the TE elements. The percentage of retained loci varied widely in the different datasets. The amount of loci shared by at least 90% of the individuals in both wolf datasets was around 25%, while in the cervid datasets it was around 4%. The low percentage of shared loci in the cervid datasets might be associated with the variable levels of DNA degradation and the distribution of the targeted TE in cervid genomes. BOV2A, the TE targeted in the cervid dataset, is a SINE element that is widely distributed in the genomes of ruminants (e.g. deer) (Nilsson et al., 2012; Onami, Nikaido, Mannen, & Okada, 2007). The high abundance of the element combined with the reduced effort on sequencing could be driving the allelic dropout. A study on sika deer using ddRAD obtained 7,576,300 candidates of the ddRAD loci from all individuals (Ba et al., 2017). After data filtering, 4% of those loci were shared by more than 75% of the individuals. Our BOV2A_CE dataset yielded similar results of shared loci with a 90% cut-off. In terms of raw reads and SNP numbers, Ba et al. (2017); sequenced ca. 34.5 million PE reads per sample, which resulted in 96,000 SNPs in the loci that were shared by more than 75% of the individuals. In comparison, we sequenced an average of 850,000 PE reads per deer sample, which yielded approximately 115,000 SNPs in the loci that were shared by at least 90% of the individuals. In this way, despite the need for adjustment in the sequencing effort, our results are comparable to similar studies.

### Length of flanking region

Available RRL methods produce loci with variable length, depending on the cutting enzyme and the chosen sequencing technology (Andrews et al., 2016). Currently, only the latest RADseq version can sequence up to 700 bp fragments (Nelson & Cresko, 2017). Our method has allowed us to sequence an average of up to 650 bp in wolf LINE, 600 bp in wolf SINE, 500 bp in deer BOV2A, and 650 bp in rat L1, with variable depth of coverage (Figure 3). Length variation is correlated with the fragment length used for library build. In this way, if the aim of the study requires a great extension into the flanking region, it would be crucial to take into consideration the fragmentation level of the starting material.

### Ease of adjustment to other taxa

In this study, we have applied MobiSeq to three different mammal groups, focusing on SINE and LINE retrotransposons. However, other genomic mobile elements could be targeted. Designing the TE-target primer requires some major considerations to be taken into account. The targeted elements need to present a conserved region for primer annealing. Many of these mobile genomic elements present terminal repetitive sequences, such as Long Terminal Repeat elements (Mourier & Willerslev, 2009). If the element of choice is characterized by terminal repetitive motifs, the amount of information in the flanking region would be reduced. Another important consideration is the distribution of the element in the genome, as it will have an impact on depth of coverage, as well as in the amount of information that can be used to performed evolutionary analyses. However, there is no taxonomic limitation for the method and, in theory, it should be applicable to most eukaryotes (Chénais et al., 2012).

Despite the benefits of MobiSeq, we caution there are some aspects that need further exploration:

1. Here, we focused on the use of the flanking region for variant calling. Another source of genomic information would be to look at the presence/absence of the loci in different samples. The SINE and LINE elements targeted in this study present a copy-and-paste mechanism of replication. Once inserted in a new genomic region, their genetic signature will be present over evolutionary relevant timescales. This information could be used in cluster analysis or presence/absence trees. However, to perform these analyses, we would need to either target elements with reduced copy number, or increase the sequencing effort, in order to mitigate the effect of allelic dropout.
2. One of the limitations of traditional RRL methods is its applicability to non-invasive samples, such as fecal samples. The high proportion of non-host DNA would affect the sequencing efficiency. Chiou and Bergey (2018) suggested a methylation-based enrichment of fecal DNA extracts prior to RRL (RADseq, in their case), as a way to reduce the non-host DNA. As MobiSeq relies on TE-target species specific primers, the presence of non-host DNA should not reduce the efficiency of the method. If MobiSeq can be implemented in non-invasive samples to generate genomic information, it will have a broad applicability in conservation genomics and wildlife forensics at a reduced cost.
3. Our BOV2A_all dataset included specimens from *Cervus* sp., as well as two fallow deer, which belong to *Dama*, a different Cervidae genus (Gilbert et al., 2006). Despite their phylogenetic distance, the same TE-target primer allowed us to sequence shared loci and enabled variant calling. Our results indicate that MobiSeq could also be applied to investigate relationships between phylogenetically distant species.

Our study presented MobiSeq, a RRL method that uses TEs as anchor for sequencing extension into the flanking region of these mobile elements. MobiSeq generates high numbers of comparable loci and SNPs across samples. We also demonstrate the ability of the method to sequence up 650 bp of the flanking region, and its adaptability to target different genome mobile elements. This makes MobiSeq a good alternative to other RRL protocols to perform evolutionary inferences.

## Acknowledgements

We are grateful to all the people and institutions that have provided samples for this study, specifically: Department of Environment Nunavut, Environment and Natural Resources Northwest Territories, and Lindsey Carmichael and David Coltman at University of Alberta (wolf samples); Frank Zachos (Natural History Museum in Vienna), Meirav Meiri (Tel-Aviv University), Adrian Lister and Ian Barnes (Natural History Museum of London), and Kristian Murphy Gregersen (Natural History Museum of Denmark) (deer material). We also thank Lasse Vinner for experimental methodology discussions. Maria Asplund for discussion on data analysis in the early stages of the project. The Danish Advance Technology Foundation. S.G. was funded by EU Marie Słodowska-Curie grant 655732 (Wherewolf).

## Author contributions

A.J.H., T.M, M.T.P.G. and M.B conceived the ideas and designed methodology. D.A.P. and A.A.N. conducted the laboratory work in the initial phases of method development. C.C. and A.R.I. optimized, implemented the laboratory method, and conducted laboratory work in the rat and deer datasets. M.H.S.S. performed the sampling and curation of the wolf material. L.B.P. and C.N.N. conducted laboratory work in the wolf dataset. M. F. R. performed rat dissection and tissue sampling. S.G., T.R., T.M., and A.R.I. conducted data analyses. Z.L. and B.P. generated and assembled the red deer genome. A.R.I. wrote the manuscript with input from all the authors. All authors read the final draft and approved it for publication.

## Data Accessibility

The data will be accessible at the Electronic Research Data Archive at the University of Copenhagen UCPH ERDA) upon acceptance for publication. All the scripts used for processing the data and generating the plots included in the main and supplementary material are available on GitHub at https://github.com/shyamsg/MobiSeq.

